# Cortisol-induced signatures of stress in the fish microbiome

**DOI:** 10.1101/826503

**Authors:** Tamsyn M. Uren Webster, Deiene Rodriguez-Barreto, Sofia Consuegra, Carlos Garcia de Leaniz

**Affiliations:** Centre for Sustainable Aquatic Research, College of Science, Swansea University, Swansea, SA2 8PP, UK

**Keywords:** Stress response, Microbiota, Glucocorticoid, *Salmo salar*, Lactic acid bacteria

## Abstract

Stress experienced in intensive aquaculture can compromise fish growth, condition and immunity. Microbiome disruption may contribute to these adverse health effects, but little is known about how stress affects fish microbial communities. Here, we specifically examined the effects of stress-induced cortisol production on the fish microbiome. We exposed juvenile Atlantic salmon to a mild confinement stressor for two weeks. We then measured cortisol in the plasma, skin-mucus and faeces, and characterised the skin and faecal microbiome. Faecal and skin cortisol concentrations increased in fish exposed to confinement stress, and were positively correlated with plasma cortisol. Elevated faecal cortisol was associated with pronounced changes in the diversity and structure of the faecal microbiome. In particular, we identified a marked decline in probiotic Lactobacillales (*Carnobacterium* sp.) and an increase in pro-inflammatory and pathogenic taxa within the classes Clostridia and Gammaproteobacteria. In contrast, skin-mucus cortisol concentrations were lower and not associated with any detectable changes in the skin microbiome. Our results demonstrate that cortisol disrupts the gut microbiome, which may, in turn, contribute to the adverse effects of stress on fish health. They also highlight the value of using non-invasive faecal samples to monitor stress, including simultaneous determination of cortisol and stress-responsive bacteria.

## Introduction

Aquaculture is the fastest growing food producing sector, and plays an increasingly important role in global food security in the face of a growing human population, depletion of capture fisheries and climate change (Teletchea & Fontaine 2014). However, the intensification of aquaculture is often associated with an increase in stress, for example due to crowding and handling, which can impact animal health and welfare, and threaten aquaculture sustainability (Iwama et al. 2011). Stress response in fish includes activation of the hypothalamus-pituitary-interrenal (HPI) axis, culminating in the release of glucocorticoids from interrenal cells located in the head kidney (Barton 2002). As for mammals, cortisol is the predominant glucocorticoid released as part of the primary stress response, and is critical for mediating adaptive metabolic, physiological and behavioural adjustments (Schreck & Tort 2016). However, prolonged elevation of cortisol is associated with adverse health effects. In particular, stress-mediated impairment of immune function has been widely described in cultured fish and has been associated with reduced disease resistance, which is of critical importance for aquaculture sustainability (Ellison et al. 2018; Uren Webster et al. 2018b; Yada & Tort 2016).

Recent research has revealed the diverse influence of microbiota and their metabolites on many aspects of host health and fitness, including digestion and nutrient uptake, metabolism and immune development (Hooper et al. 2012; Rea et al. 2016). In mammals, stress is well known to disrupt the diversity, structure and function of the microbiome which, in turn, has been associated with long term health effects in the host, including metabolic and immune impairment, and a range of diseases (Foster et al. 2017; Tetel et al. 2018). The mechanisms by which stress impacts the microbiome are complex, and not fully understood. Elevated plasma cortisol, resulting from social or psychological stress, has been associated with alterations in the structure and/or diversity of the mammalian faecal or oral microbiome, including the abundance of lactic acid bacteria and opportunistic pathogens (Galley et al. 2014; Jasarevic et al. 2015; Mudd et al. 2017). Direct administration of glucocorticoids has also been shown to exert stimulatory and inhibitory effects on specific microbial taxa, together with wider effects on host metabolism in some cases (Huang et al. 2015; Jentsch et al. 2013; Petrosus et al. 2018; Wu et al. 2018). Additionally, while host stress response influences the microbiome, the microbiome can also influence host stress response. Microbiota and their metabolites are known to exert effects throughout the mammalian hypothalamus-pituitary-adrenal (HPA) axis, influencing glucocorticoid synthesis, release and signalling pathways (Burokas et al. 2017; de Weerth 2017; Simard et al. 2014; Vodicka et al. 2018).

Disruption of the microbiome is likely to represent an important mechanism by which stress affects fish health, welfare and performance in aquaculture. There is some evidence that environmental and social stressors disrupt microbial communities associated with the fish gut and skin (Boutin et al. 2013; Sylvain et al. 2016; Uren Webster et al. 2019; Zha et al. 2018), however, a potential role of cortisol in mediating these effects is unknown. We hypothesised that stress-induced cortisol production would directly disrupt the fish microbiome. To test this, we examined the effects of cortisol on the skin and gut microbiome of juvenile Atlantic salmon, following confinement, a mild aquaculture-relevant stressor.

## Methods

### Stress experiment

Prior to the start of the experiment 0+ Atlantic salmon fry (mean mass 3.92 ± 0.11 g; fork length 7.46 ± 0.07 cm), were housed in stock tanks (80 L) supplied with a constant flow of aerated, de-chlorinated tap water in a recirculation system with a temperature of 15°C ± 0.5 °C and photoperiod of 12L:12D. Water oxygen saturation was maintained above 90%, and ammonia (<0.02 mg/L), nitrite (<0.01 mg/L), nitrate (<15 mg/L) and pH (7.5 ± 0.2) were maintained within the optimal range for the species. Fish were fed with a commercial salmon feed (Skretting Nutra Parr) at a rate of 3% body weight per day.

Experimental fish were assigned at random to the control and confinement-stress treatment groups, with three replicate 20 L tanks per group, each containing 28 fish. Confinement stress consisted of slowly lowering the water volume in each tank (via draining) from 20 L to 5 L for one hour, and this was repeated every day at the same time (1100 hrs) for two weeks. All other husbandry conditions were as before. At the end of the experiment, fish were euthanised via an overdose of anaesthetic (Phenoxyethanol; 0.5 mg/L), followed by destruction of the brain according to UK Home Office regulations. The fish were measured (fork length), weighed (wet weight) and Fulton’s condition factor was calculated as a measure of body condition. Blood samples were collected from the caudal vein using heparinised capillary tubes, centrifuged (5 min, 5000 x *g*) and the plasma supernatant removed and stored at −80 °C prior to cortisol analysis. For each fish, a sample of skin-associated mucus for microbiome analysis was collected by swabbing the left side lateral line five times using Epicentre Catch-All™ Sample Collection Swabs (Cambio, Cambridge, UK). A sample of skin-associated mucus for cortisol analysis was collected by scraping mucus from a 2 cm^2^ region of skin on the left-hand side of the fish, above the lateral line between the head and dorsal fin, using a scalpel blade. Faecal samples were collected from each fish by gently pressing along the length of the abdomen and collecting expelled faeces, which were then split evenly between samples for cortisol and microbiome analysis. All skin-mucus and faecal samples were directly frozen at −80 °C prior to analysis.

### Cortisol measurement

Quantification of cortisol concentration in plasma, skin mucus and faecal samples was performed using the DetectX Cortisol Enzyme Immunoassay Kit (Arbor Assays, Michigan, USA), according to the manufacturer’s recommendations. Briefly, plasma samples were first pre-treated with dissociation reagent then diluted in assay buffer (1:50 final dilution) before cortisol measurement. Faecal samples were suspended in 100 µl ethanol, vortexed for 30 minutes, centrifuged (5 min, 5000 x *g*), then the supernatant was then diluted in assay buffer (1:20) before cortisol measurement. Skin-mucus samples were suspended in 100 µl 1M Tris-HCl, vortexed for 30 minutes and centrifuged (5 min, 5000 x *g*), then the supernatant was used directly in the assay without dilution.

Cortisol concentration was measured in the plasma, faeces and skin-mucus for a total of 60 individual fish (40 stressed fish and 20 controls; distributed evenly amongst replicate tanks). Each of these 180 samples was analysed in duplicate, across five 96-well plates. Cortisol concentration was calculated based on a standard curve run on each plate, and adjusted for dilution factor and initial sample volume (plasma) or weight (skin-mucus/faeces). Inter-assay variability, measured as the coefficient of variation (CV%) of four repeats across the five plates, was 4.62% and the lower limit of detection was 76.4 pg/ml. We removed one outlier of faecal cortisol (47.3 ng/g) from the stressed group (most likely resulting from an error in sample preparation) using Tukey’s 1.5*IQR method, as it was 3.4x higher than the mean value (13.76 ng/g) and 1.6x higher than the next highest value in this group (29.1 ng/g).

### 16S rRNA amplicon sequencing

16S rRNA amplicon sequencing was performed using the faecal and skin mucus samples for the same 60 individual fish for which cortisol quantification was performed (40 stressed and 20 controls), as described previously (Uren Webster et al. 2018a). Briefly, DNA was extracted from all samples using the MoBio PowerSoil® DNA Isolation Kit (Qiagen) according to the manufacturer’s instructions. Libraries were prepared amplifying the V4 hypervariable region of the bacterial 16S gene using the primers 341F and 785R, and sequenced across two lanes of an Illumina MiSeq. Raw sequence reads were quality filtered using Trimmomatic (Bolger et al. 2014), before analysis with mothur v1.39 (Kozich et al. 2013). Concatenated reads were aligned to the Silva seed reference database (version 128) (Quast et al. 2013), chimeric reads were removed using UCHIME (Edgar et al. 2011), and Bacteria and Archaea contigs were classified using the Silva reference taxonomy. Contigs were clustered into operational taxonomic units (OTUs) using mothur, based on 97% sequence similarity. Singleton OTUs were removed from the dataset then all faecal samples were subsampled to an equal depth of 19,724 reads and all skin samples were subsampled to 10,133 reads. Measures of alpha diversity (Chao1 richness and Shannon diversity) for each sample were calculated in mothur.

### Statistical analysis

All statistical analysis was performed in R v3.5.0. Firstly, we assessed whether non-invasive measurements of faecal and skin cortisol were indicative of cortisol in blood plasma by calculating the Pearson correlation coefficient. We then employed linear mixed effects models (LMM) using the *lme4* package to examine the effects of confinement stress and fish size on measured cortisol values in plasma, skin and faeces, using tank identity as a random factor. We also used linear mixed effects models to examine the effects of confinement stress, fish size and measured faeces/skin cortisol on faces/skin microbial alpha diversity (Chao1 richness and Shannon diversity), including tank as a random factor. We used fish length as a covariate to control for size effects as it had a lower coefficient of variation (CV = 0.067) than fish mass (CV = 0.216). To achieve model simplification, we started with a model with all main effects and selected the model with the lowest AIC value via backward selection using the *step* and *drop1* functions and the *lmerTest* package (Kuznetsova et al. 2017). A minimal adequate model was then refitted via Restricted Maximum Likelihood, or as a linear model when the random component (tank identity) did not improve model fit compared to the fixed effects only model, as indicated by the Likelihood Ratio Test (LRT). We used the *VCA* package to estimate the amount of variability in cortisol due to confinement stress, tank effects, and differences among individual fish.

Analysis of microbial community structure (beta diversity) was performed within the Vegan package in R (Oksanen et al. 2017), using the Bray-Curtis dissimilarity index. Non-metric multidimensional scaling (NMDS) ordination of Bray-Curtis distances were visualised, including measured cortisol concentration as an environmental vector. Multivariate statistical analysis of microbial community separation in the faecal and skin samples was performed by PERMANOVA using *Adonis* in the *Vegan* package with confinement stress and measured faecal/skin cortisol as predictors. Statistical analysis of OTU abundance was performed using DeSeq2 (Love et al. 2014). The effect of confinement stress and faecal/skin cortisol on relative abundance of faecal/skin OTUs was tested using a multifactorial design. Within the DeSeq model, low coverage OTUs were independently filtered to optimise power for identification of differentially abundant OTUs at a threshold of alpha=0.05. Outlier detection and moderation of OTU level dispersion estimates were performed using default settings, and OTUs were considered significantly differentially abundant at FDR <0.05.

## Results

### Relation between plasma cortisol and non-invasive measures of cortisol in faeces and skin

Cortisol concentrations ranged from 2.9 to 65.8 ng/ml in blood plasma, 3.6 to 29.1 ng/g in faeces, and 0.14 to 9.45 ng/g in skin mucus across all samples. Significant positive correlations were found between plasma and faecal cortisol (Pearson’s *r*_*56*_=0.615, *P* < *0.001*), between plasma and skin cortisol (*r*_*56*_=0.289, *P* = *0.028*), and between faecal and skin cortisol (*r_57_*=0.422, *P* < *0.001;* Figure 1a). Variance component analysis indicated that 82-85% of the variation in cortisol was due to variation between individuals, and 0-18% was due to variation between tanks.

**Figure 1.**
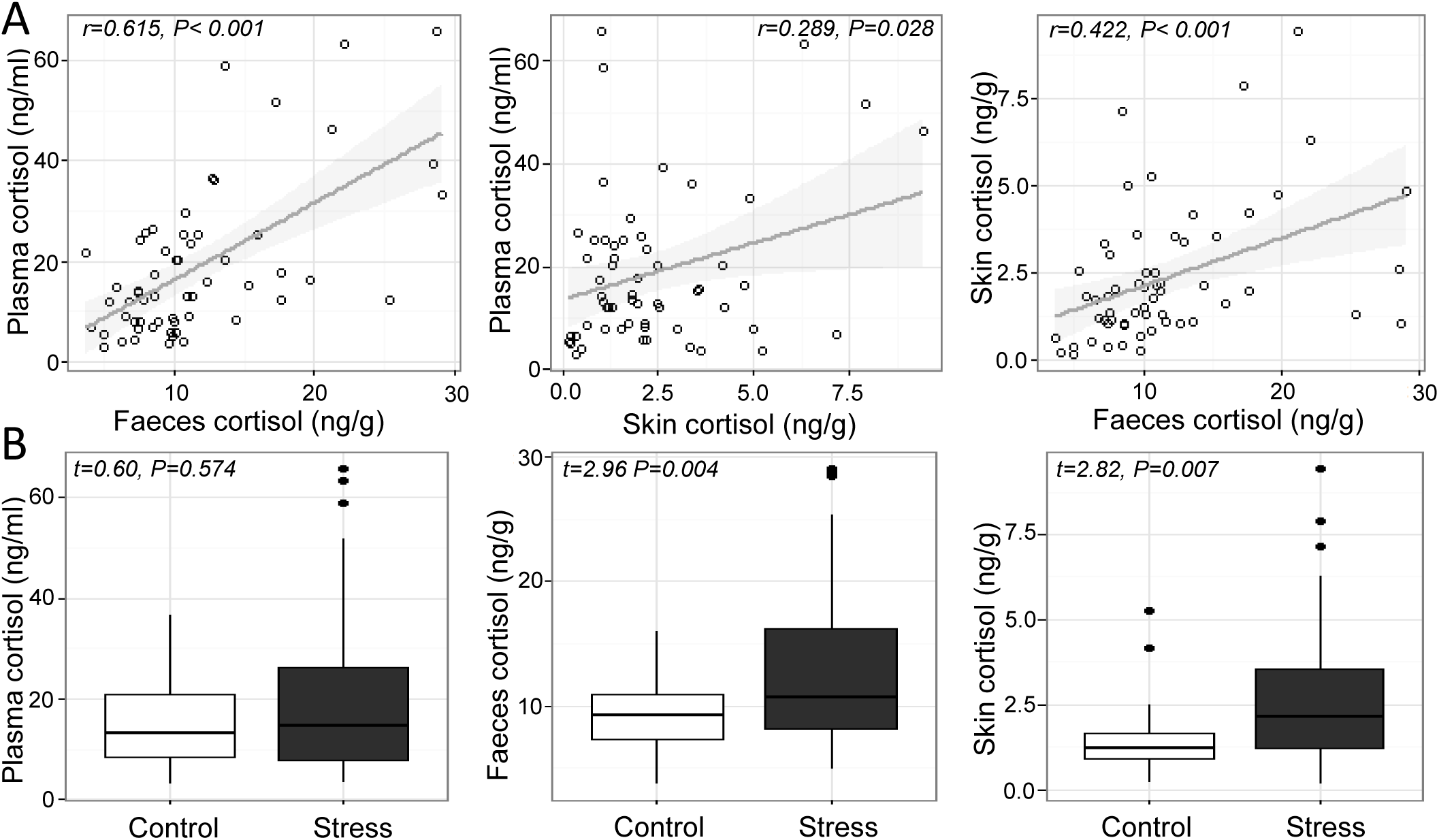
A) Relation between measured cortisol in the plasma, skin and faeces across individual fish, and B) measured cortisol in the plasma, faeces and skin of Atlantic salmon exposed to confinement stress compared to control fish.

### Effects of confinement stress on cortisol

There was a significant increase in cortisol in the faeces and skin of stressed fish compared to unstressed controls (Welch two sample t-test; faeces, *t*_56.766_ = 2.955, *P* = 0.004; skin, *t*_52.917_ = 2.819, *P* = 0.007), but not in blood plasma (LMM stress effect, *t*_4.785_ = 0.603, *P* = 0.574; Figure 1b), which showed a significant tank effect (LRT ^2^ = 6.01, *P* = 0.014). There was no association between cortisol and the length, weight or body condition of fish at the end of the experiment (*P* >0.2 in all cases).

### Effects of cortisol on microbial diversity

Faecal cortisol was negatively correlated with faecal Chao1 microbial richness (Chao1 Cortisol estimate: −79.17 ± 32.33, *t*_1,57_ = −2.449, *P* = 0.017), but positively correlated with Shannon diversity (Shannon Cortisol estimate: 0.08 ± 0.024, t_1,57_ = 3.536, *P* < 0.001; Figure 2). There was no effect of confinement stress, fish size or tank identity on faecal microbial diversity beyond that accounted by an increase in cortisol (P>0.4 in all cases). For the skin, there was no significant effect of confinement stress, skin cortisol, or fish size on skin microbial Chao1 richness of Shannon diversity (P>0.1 in all cases), although there were significant tank effects for skin Chao1 richness (LRT *χ*^2^ = 15.53, *P* < 0.001).

**Figure 2.**
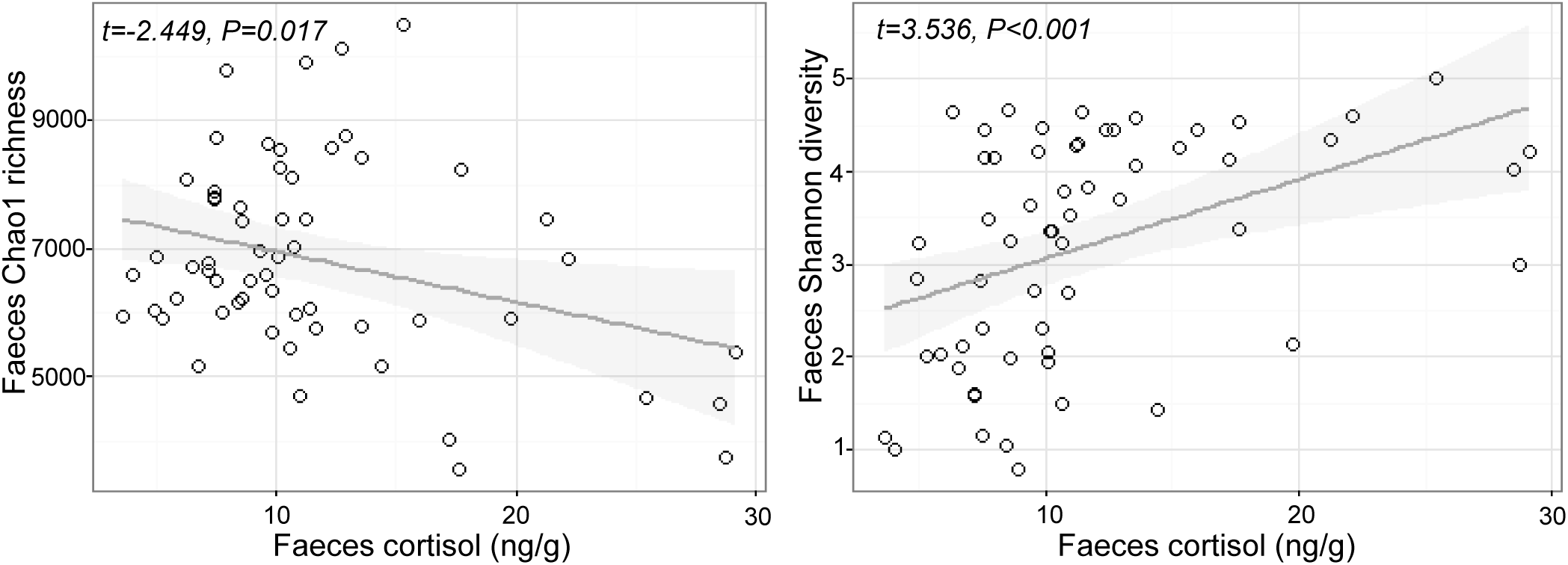
Relationship between measured cortisol and microbial alpha diversity (Chao1 richness and Shannon diversity) in the faeces.

Microbial community structural diversity was performed based on the Bray-Curtis dissimilarity metric, and visualised using NMDS analysis (Figure S1). As for alpha diversity, there was a significant effect of faecal cortisol on faecal microbiome beta diversity (Cortisol: F_1,56_ = 9.525, *P* = 0.001), but the confinement stress had no additional effect beyond that caused by an increase in cortisol (Stress F_1,56_ = 2.487, P = 0.064). For the skin microbiome there was no detectable effect of stress or skin cortisol concentration on beta diversity (Stress: F_1,53_ = 1.263, P=0.151, Cortisol: F_1,53_ = 0.960, P=0.475).

### Effects of cortisol on microbial composition

The effects of stress and cortisol concentration on OTU relative abundance was investigated using DeSeq2. For the faecal microbiome, the abundance of 44 OTUs (27 increased, 17 decreased) were significantly associated with faecal cortisol concentration, but only one OTU (*Vagacoccus* sp.) was significantly elevated in the confinement stress group independently of cortisol (Figure 3; Table S1). Strikingly, of the 17 OTUs which were negatively associated with cortisol concentration, the vast majority (15) were classified as belonging to the genus *Carnobacterium* sp. in the order Lactobacillales, including the most abundant OTU overall. Of the OTUs that were positively associated with faecal cortisol concentration, 10 (37%) were from the class Gammaproteobacteria and, notably, two highly abundant OTUs from the family Clostridiaceae were also increased. In contrast, for the skin microbiome, no OTUs were significantly associated with either confinement stress or measured skin cortisol concentration.

**Figure 3.**
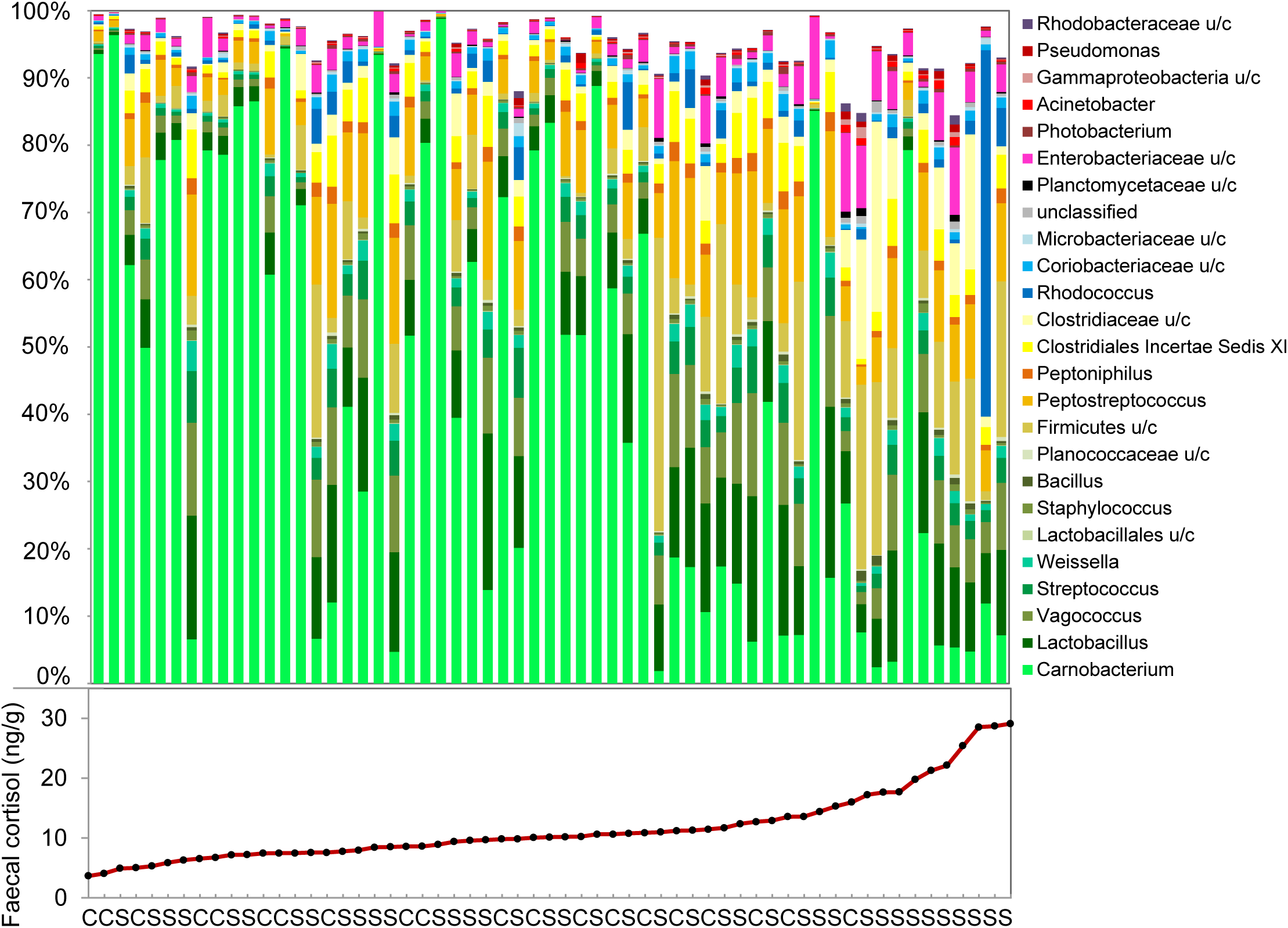
Genus-level composition of the faecal microbiome, and measured faecal cortisol concentrations in individual fish. (C: control, S: confinement stress).

## Discussion

Our study indicates that the cortisol stress response modulates the intestinal, but not the skin, microbiome of juvenile Atlantic salmon. Although the cortisol stress response to confinement was variable amongst individuals, we identified a very distinctive relationship between cortisol and the diversity and structure of the faeces microbiome, including a clear inhibition of lactic acid-producing bacteria and the promotion of pro-inflammatory and pathogenic taxa.

Exposure to a routine, aquaculture-relevant stressor (confinement) increased the cortisol concentration in the skin and faeces of juvenile salmon compared to control fish, although there was considerable variation among individuals. This variation is consistent with the existence of low and high cortisol response fish (e.g. Pottinger & Carrick 1999; Samaras et al. 2016). We also identified a positive association between plasma and faecal cortisol, and, to a lesser extent, between plasma and skin-mucus cortisol. Plasma cortisol is typically used to measure the stress response in fish, but blood sampling is invasive and may require terminal sampling in the case of small fish (Sadoul & Geffroy 2019). Plasma cortisol concentration is also known to be influenced by acute spikes in glucocorticoid production, for example caused by handling stress prior to sampling, which may mask underlying stress levels (Bertotto et al. 2010). Our results add to that of other recent studies suggesting that faecal and skin sampling provide non-destructive alternatives to measuring plasma cortisol in Atlantic salmon (Bertotto et al. 2010; Cao et al. 2017; De Mercado et al. 2018; Lupica & Turner Jr 2009), which can also be linked directly to microbiome analysis.

Across all fish, we identified a strong association between cortisol in the faeces and both alpha and beta measures of gut microbiome diversity. Faecal cortisol was negatively associated with Chao1 richness, but positively associated with Shannon diversity, suggesting that there were fewer, but more evenly distributed, bacterial taxa in the intestine of stressed fish. This is likely to reflect an inhibitory effect of cortisol on dominant OTUs normally present in non-stressed individuals. In particular, there was a striking decline in *Carnobacterium* sp. with increasing levels of faecal cortisol, including the most abundant OTU in non-stressed fish, together with +10 other OTUs assigned to this genus. *Carnobacterium* (order Lactobacillales, class Bacilli, phylum Firmicutes) is a genus of facultatively anaerobic, cold tolerant lactic acid bacteria comprising +12 species (Pikuta & Hoover 2014). This genus, particularly *C. pisciola, C. divergens* and *C. inhibens*, is commonly found in the intestinal communities of healthy fish, including Atlantic salmon (Ringø et al. 2001). *Carnobacterium* sp. are also widely used as probiotics in aquaculture, due to their beneficial effects on gut health and fish performance, and their ability to inhibit the growth of several common fish pathogens (Ringø 2008; Ringø et al. 2001).

Individuals that displayed a high cortisol response to confinement stress had a distinct faecal microbiome, that was very different from that of non-responsive fish, or from control fish that had low baseline cortisol levels. Alongside a marked decline in *Carnobacterium* sp., this structural change was characterised by a notable increase in the relative abundance of two Clostridiaceae OTUs. This family (class Clostridia, phylum Firmicutes) is commonly found in the gut of healthy mammals and fish, but also includes a number of opportunistic pathogens. An increased abundance of Clostridiaceae has been associated with microbial dysbiosis, intestinal inflammation and gastrointestinal diseases (Lopetuso et al. 2013; Muñiz Pedrogo et al. 2018). Several OTUs within the class Gammaproteobacteria, including two *Yersinia* sp., *Pseuodomonas* sp., *Acinetobacter* sp. and *Aeromonas* sp, were also particularly abundant in fish that had high levels of faecal cortisol. These genera include a range of opportunistic fish pathogens (Austin & Austin 2007), and tend to increase following exposure to different types of environmental stress in fish (Boutin et al. 2013; Uren Webster et al. 2019), suggesting they may represent a common signature of stress exposure. In mammals, experimental administration of cortisol results in a similar reduction in probiotic lactic acid-producing bacteria and an increase in pro-inflammatory microbiota (Huang et al. 2015; Petrosus et al. 2018; Wu et al. 2018), suggesting that these bacteria taxa could represent useful biomarkers of stress across vertebrates.

Our results demonstrate how an increase in cortisol can affect the diversity and structure of the salmon gut microbiome, which may in turn contribute to the adverse effects of stress on fish health. This is consistent with the results of previous studies that experimentally administered glucocorticoids to rats, mice and pigs (Huang et al. 2015; Petrosus et al. 2018; Wu et al. 2018). However, it is not clear exactly how cortisol may affect different taxa within complex host-associated microbial communities. Potential inhibitory mechanisms could include direct toxicity, metabolic impairment, disruption of ion-regulation, endocrine signalling or nutrient depletion, similar to the effects of other chemical or physical stressors (Harms et al. 2016; Weber et al. 2014). At the same time, taxa more tolerant of cortisol may flourish in the absence of previous niche completion (Hibbing et al. 2010), and cortisol is also known to specifically promote the growth of certain oral pathogens *in vitro* (Jentsch et al. 2013). However, the overall relationship between stress response, cortisol and the microbiome is complex. The microbiome may be influenced by other stress hormones, by interactions amongst microbes or with the host immune system, while microbiota and/or their metabolites are also well known to influence host stress response signalling (Burokas et al. 2017; de Weerth 2017; Simard et al. 2014; Vodicka et al. 2018).

The impacts of cortisol on the microbiome are also likely to depend on the nature of the microbial community, and the stress response. In contrast to the faecal microbiome, we found no significant effects of confinement stress or skin cortisol on the diversity or structure of the skin microbiome. It is possible that skin-associated communities, which are dominated by Proteobacteria with much lower levels of Lactobacilliales, are less sensitive to cortisol than faecal microbiota. On the other hand, cortisol concentrations in the skin mucus were much lower than those measured in faecal samples, which may also help explain the lack of observed effects of skin cortisol on the skin microbiome.

To conclude, our study shows, for the first time, that an increase in stress-related cortisol alters the fish intestinal microbiome, notably by decreasing the abundance of *Carnobacterium*, a Lactobacilliales commonly used as a probiotic in aquaculture, and increasing pro-inflammatory and opportunistic bacterial pathogens. Given the fundamental influence of microbiota and their metabolites on many aspects of host health, this suggests that cortisol-mediated disruption of the intestinal microbiome is likely to contribute to the adverse effects of stress on immune function and disease resistance. These results have important implications for health and welfare of fish exposed to environmental stress, and more broadly, to research on stress-related diseases, such as metabolic syndrome, obesity and IBD, which have been associated with microbiome dysbiosis. Finally, our study demonstrates that both cortisol measurements and microbiome analysis can be performed simultaneously on faecal and skin samples collected non-invasively, which could represent a valuable screening tool for evaluating stress in fish.

## Acknowledgements

We thank the staff at CSAR for help with fish husbandry, Nikita Berry for assistance with sample preparation and Matthew Hitchings for performing the Illumina sequencing.

## Author contributions

TUW, DRB, CGL and SC designed the study; TUW and DRB performed the experiment; TUW and CGL analysed the data; TUW drafted the manuscript. All authors contributed to the final version of the manuscript.

## Ethics

All experimental procedures were approved by Swansea Animal Welfare and Ethical Review Body (number IP-1415-2).

## Data accessibility

All 16S rRNA sequence reads are available from the European Nucleotide Archive under accession PRJEB32276. Full metadata, and measured cortisol concentrations are available in the supporting information.

