## Supporting information for "Cortisol-induced signatures of stress in the fish microbiome"

**P2-3: Table S1.** Faecal OTUs significantly associated with faecal cortisol concentration.

**P4: Figure S1.** Structural analysis of the faecal microbiome.

**P5-6: Table S2.** Morphological and cortisol measurements for all fish.

**Table S1.** Faecal OTUs significantly associated with faecal cortisol concentration.

| OTU | Phylum | Class | Order | Family | Genus | Log2 Fold Change | FDR |
|---|---|---|---|---|---|---|---|
| Otu000001 | Firmicutes | Bacilli | Lactobacillales | Carnobacteriaceae | Carnobacterium | -0.29 | 0.00008 |
| Otu000005 | Firmicutes | Clostridia | Clostridiales | Clostridiaceae | unclassified | 0.15 | 0.00835 |
| Otu000006 | Firmicutes | Clostridia | Clostridiales | Clostridiaceae | unclassified | 0.19 | 0.02552 |
| Otu000009 | Actinobacteria | Actinobacteria | Actinomycetales | Nocardiaceae | Rhodococcus | 0.13 | 0.00651 |
| Otu000014 | Actinobacteria | Actinobacteria | Actinomycetales | Nocardiaceae | Rhodococcus | 0.11 | 0.03822 |
| Otu000018 | Proteobacteria | Gammaproteobacteria | Enterobacteriales | Enterobacteriaceae | Yersinia | 0.39 | 0.00002 |
| Otu000022 | Proteobacteria | Gammaproteobacteria | Enterobacteriales | Enterobacteriaceae | Yersinia | 0.20 | 0.02120 |
| Otu000023 | Proteobacteria | Gammaproteobacteria | Enterobacteriales | Enterobacteriaceae | Buttiauxella | -0.30 | 0.00236 |
| Otu000032 | Firmicutes | Bacilli | Lactobacillales | Lactobacillaceae | Lactobacillus | 0.07 | 0.00069 |
| Otu000053 | Proteobacteria | Gammaproteobacteria | Pseudomonadales | Pseudomonadaceae | Pseudomonas | 0.18 | 0.00580 |
| Otu000055 | Firmicutes | Bacilli | Lactobacillales | Carnobacteriaceae | Carnobacterium | -0.37 | 0.00003 |
| Otu000056 | Firmicutes | Bacilli | Bacillales | Bacillaceae | Bacillus | 0.21 | 0.00835 |
| Otu000059 | Proteobacteria | Gammaproteobacteria | Pseudomonadales | Moraxellaceae | Acinetobacter | 0.12 | 0.02122 |
| Otu000060 | Proteobacteria | Gammaproteobacteria | Aeromonadales | Aeromonadaceae | Aeromonas | 0.17 | 0.00685 |
| Otu000061 | Firmicutes | Bacilli | Lactobacillales | Carnobacteriaceae | Carnobacterium | -0.27 | 0.00134 |
| Otu000067 | Proteobacteria | Alphaproteobacteria | Rhodobacterales | Rhodobacteraceae | unclassified | 0.22 | 0.00383 |
| Otu000074 | Firmicutes | Bacilli | Bacillales | unclassified | unclassified | 0.23 | 0.01224 |
| Otu000076 | Tenericutes | Mollicutes | Mycoplasmatales | Mycoplasmataceae | Mycoplasma | 0.37 | 0.00037 |
| Otu000086 | Actinobacteria | Actinobacteria | Actinomycetales | Micrococcaceae | Micrococcus | 0.11 | 0.03692 |
| Otu000092 | Firmicutes | Bacilli | Lactobacillales | Carnobacteriaceae | Carnobacterium | -0.26 | 0.00265 |
| Otu000099 | Firmicutes | Clostridia | Clostridiales | Clostridiaceae | Clostridium | 0.21 | 0.00685 |
| Otu000100 | Proteobacteria | Gammaproteobacteria | unclassified | unclassified | unclassified | 0.38 | 0.00265 |
| Otu000103 | Firmicutes | unclassified | unclassified | unclassified | unclassified | 0.17 | 0.00215 |
| Otu000104 | Proteobacteria | Gammaproteobacteria | Pseudomonadales | Pseudomonadaceae | Pseudomonas | 0.14 | 0.00822 |
| Otu000107 | Firmicutes | Bacilli | Lactobacillales | Carnobacteriaceae | Carnobacterium | -0.28 | 0.00106 |
| Otu000108 | Firmicutes | Bacilli | Lactobacillales | Carnobacteriaceae | Carnobacterium | -0.32 | 0.00404 |
| Otu000110 | Proteobacteria | Gammaproteobacteria | Pseudomonadales | Moraxellaceae | Acinetobacter | 0.16 | 0.04834 |

| | | | | | | | |
|---|---|---|---|---|---|---|---|
| Otu000111 | Firmicutes | unclassified | unclassified | unclassified | unclassified | 0.28 | 0.00265 |
| Otu000113 | Firmicutes | unclassified | unclassified | unclassified | unclassified | 0.21 | 0.02942 |
| Otu000119 | Firmicutes | Bacilli | Lactobacillales | Carnobacteriaceae | Carnobacterium | -0.29 | 0.00106 |
| Otu000124 | Firmicutes | Bacilli | Lactobacillales | Carnobacteriaceae | Carnobacterium | -0.27 | 0.00338 |
| Otu000129 | Proteobacteria | Gammaproteobacteria | Pseudomonadales | Moraxellaceae | Enhydrobacter | 0.16 | 0.03692 |
| Otu000148 | Proteobacteria | Gammaproteobacteria | Xanthomonadales | Xanthomonadaceae | Arenimonas | 0.27 | 0.02271 |
| Otu000149 | Firmicutes | Bacilli | Lactobacillales | Carnobacteriaceae | Carnobacterium | -0.26 | 0.01513 |
| Otu000153 | Firmicutes | Bacilli | Lactobacillales | Carnobacteriaceae | Carnobacterium | -0.45 | 0.00160 |
| Otu000156 | Firmicutes | Bacilli | Lactobacillales | Carnobacteriaceae | Carnobacterium | -0.33 | 0.00189 |
| Otu000160 | Firmicutes | Bacilli | Lactobacillales | Carnobacteriaceae | Carnobacterium | -0.37 | 0.00027 |
| Otu000162 | Firmicutes | Bacilli | Bacillales | Planococcaceae | Lysinibacillus | 0.12 | 0.00338 |
| Otu000173 | Firmicutes | Bacilli | Lactobacillales | Carnobacteriaceae | Carnobacterium | -0.32 | 0.00651 |
| Otu000177 | Firmicutes | Bacilli | Bacillales | Bacillaceae | Bacillus | 0.15 | 0.00785 |
| Otu000180 | Firmicutes | Bacilli | Lactobacillales | Carnobacteriaceae | Carnobacterium | 0.16 | 0.03573 |
| Otu000185 | Firmicutes | Bacilli | Lactobacillales | Carnobacteriaceae | Carnobacterium | -0.38 | 0.00106 |
| Otu000194 | Firmicutes | Bacilli | Lactobacillales | Carnobacteriaceae | Carnobacterium | -0.32 | 0.00287 |
| Otu000200 | Actinobacteria | Actinobacteria | Actinomycetales | Dietziaceae | Dietzia | -0.19 | 0.00709 |

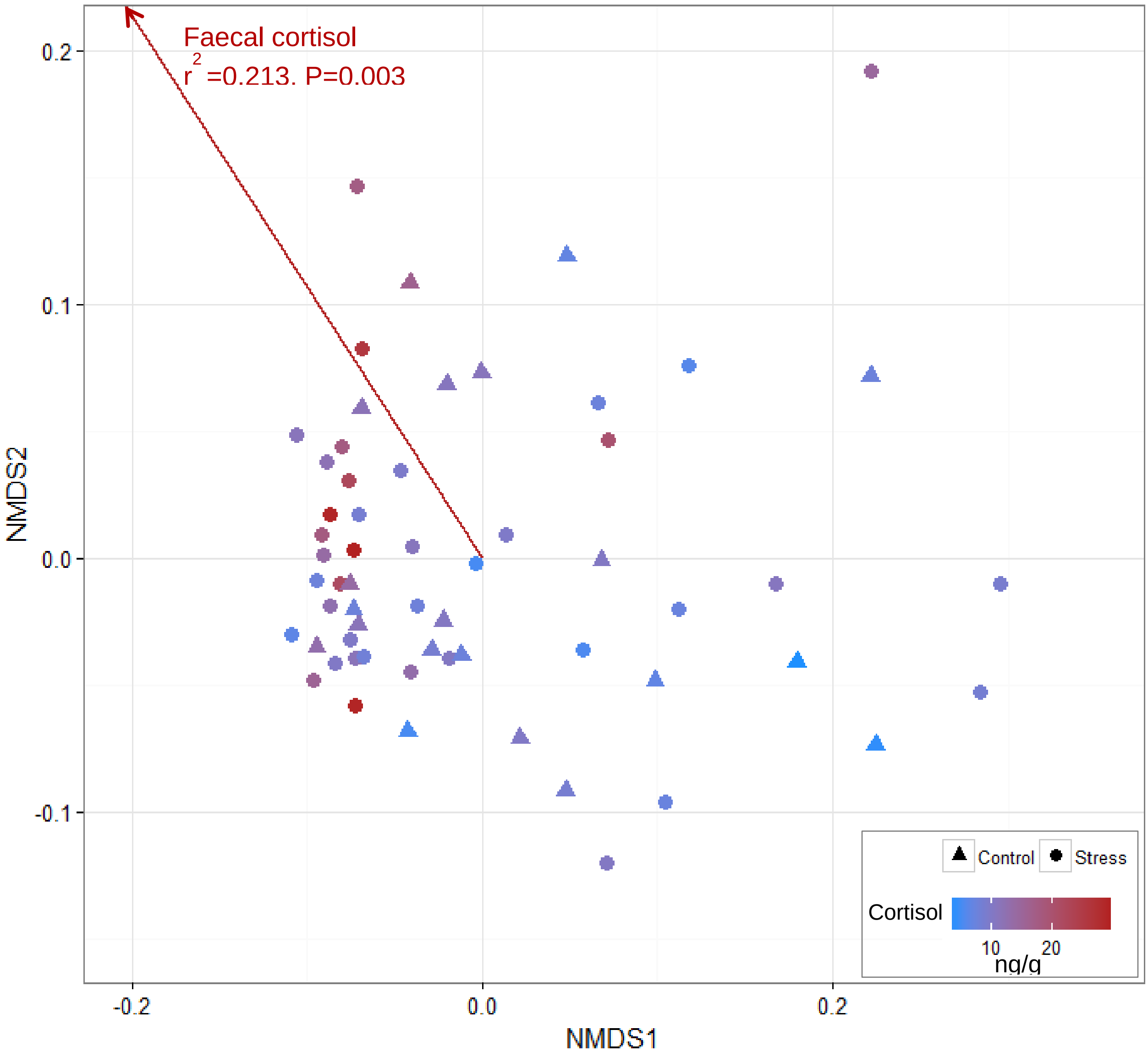


**Figure S1.** Structural analysis of the faecal microbiome. Non-metric multidimensional scaling ordination was performed based on the Bray-Curtis dissimilarity index, and correlated with measured faecal cortisol concentration. Individual points are colour-coded based on cortisol concentration.

**Table S2.** Morphological and cortisol measurements for all fish.

| Fish ID | Tank | Treatment | Faeces cortisol (ng/g) | Skin cortisol (ng/g) | Plasma cortisol (ng/ml) | Weight (g) | Fork length (cm) | Condition factor |
|---|---|---|---|---|---|---|---|---|
| 53 | Control A | Control | 3.65 | 0.63 | 21.50 | 5.35 | 8.2 | 0.97 |
| 51 | Control A | Control | 4.06 | 0.21 | 6.66 | 3.66 | 7.4 | 0.90 |
| 21 | Control B | Control | 5.00 | 0.37 | 2.97 | 5.96 | 8.0 | 1.16 |
| 26 | Control B | Control | 6.55 | 1.73 | 8.86 | 4.15 | 7.4 | 1.02 |
| 82 | Control C | Control | 6.73 | 1.19 | 11.99 | 5.58 | 8.5 | 0.91 |
| 54 | Control A | Control | 7.44 | 0.33 | 6.31 | 4.33 | 7.9 | 0.88 |
| 30 | Control B | Control | 7.45 | 1.03 | 14.08 | 4.99 | 7.9 | 1.01 |
| 28 | Control B | Control | 7.55 | 1.36 | 24.14 | 4.60 | 7.8 | 0.97 |
| 59 | Control A | Control | 8.57 | 1.04 | 13.06 | 3.27 | 7.1 | 0.91 |
| 58 | Control A | Control | 8.61 | 0.97 | 17.28 | 3.94 | 7.8 | 0.83 |
| 57 | Control A | Control | 9.82 | 0.65 | 8.75 | 3.77 | 7.4 | 0.93 |
| 83 | Control C | Control | 10.06 | 1.52 | 7.91 | 4.84 | 8.2 | 0.88 |
| 56 | Control A | Control | 10.23 | 1.3 | 20.36 | 3.48 | 7.3 | 0.89 |
| 90 | Control C | Control | 10.62 | 5.24 | 3.76 | 3.88 | 7.5 | 0.92 |
| 86 | Control C | Control | 10.86 | 2.47 | 12.98 | 6.23 | 8.5 | 1.01 |
| 25 | Control B | Control | 11.20 | 2.18 | 23.53 | 5.40 | 8.6 | 0.85 |
| 87 | Control C | Control | 11.43 | 1.27 | NA | 3.60 | 7.0 | 1.05 |
| 27 | Control B | Control | 12.69 | 1.05 | 36.65 | 3.86 | 8.0 | 0.75 |
| 84 | Control C | Control | 13.56 | 4.17 | 20.11 | 3.43 | 7.2 | 0.92 |
| 88 | Control C | Control | 15.97 | 1.6 | 25.22 | 3.21 | 7.0 | 0.93 |
| 32 | stress B | Stress | 4.92 | 0.14 | 5.38 | 6.02 | 8.5 | 0.98 |
| 9 | stress A | Stress | 5.29 | 2.53 | 11.99 | 6.54 | 8.9 | 0.93 |
| 38 | stress B | Stress | 5.86 | 1.83 | 14.69 | 4.27 | 7.9 | 0.87 |
| 61 | stress C | Stress | 6.30 | 0.48 | 3.96 | 3.85 | 7.6 | 0.88 |
| 62 | stress C | Stress | 7.18 | 1.11 | 8.00 | 4.83 | 7.8 | 1.02 |
| 6 | stress A | Stress | 7.20 | 3.33 | 4.45 | 4.01 | 7.7 | 0.88 |
| 63 | stress C | Stress | 7.45 | 1.81 | 13.64 | 7.39 | 9.0 | 1.01 |
| 10 | stress A | Stress | 7.55 | 3.02 | 7.91 | 4.90 | 8.0 | 0.96 |
| 7 | stress A | Stress | 7.76 | 1.14 | 12.16 | 5.17 | 8.3 | 0.90 |
| 68 | stress C | Stress | 7.95 | 2.03 | 25.81 | 3.37 | 7.1 | 0.94 |
| 34 | stress B | Stress | 8.43 | 0.38 | 26.52 | 4.34 | 7.7 | 0.95 |
| 8 | stress A | Stress | 8.50 | 7.14 | 6.77 | 3.46 | 7.5 | 0.82 |
| 66 | stress C | Stress | 8.89 | 4.98 | 7.90 | 4.58 | 7.9 | 0.93 |
| 45 | stress B | Stress | 9.37 | 1.36 | 21.84 | 3.34 | 7.1 | 0.93 |
| 4 | stress A | Stress | 9.57 | 3.59 | 3.49 | 4.53 | 7.8 | 0.96 |
| 3 | stress A | Stress | 9.68 | 2.19 | 5.70 | 5.81 | 8.4 | 0.98 |
| 31 | stress B | Stress | 9.83 | 0.23 | 5.06 | 5.06 | 8.1 | 0.95 |
| 71 | stress C | Stress | 10.12 | 2.08 | 5.70 | 3.37 | 7.1 | 0.94 |
| 11 | stress A | Stress | 10.17 | 2.47 | 20.11 | 4.16 | 7.6 | 0.95 |
| 69 | stress C | Stress | 10.61 | 0.8 | 25.13 | 3.76 | 7.2 | 1.01 |
| 39 | stress B | Stress | 10.75 | 1.75 | 29.61 | 4.96 | 8.4 | 0.84 |
| 5 | stress A | Stress | 10.97 | 2.13 | 8.98 | 5.57 | 8.2 | 1.01 |

| | | | | | | | | |
|---|---|---|---|---|---|---|---|---|
| 12 | stress A | Stress | 11.27 | 1.98 | 12.98 | 4.42 | 7.7 | 0.97 |
| 14 | stress A | Stress | 11.67 | 1.09 | 25.22 | 5.30 | 8.5 | 0.86 |
| 70 | stress C | Stress | 12.35 | 3.55 | 15.81 | 3.37 | 6.6 | 1.17 |
| 64 | stress C | Stress | 12.90 | 3.36 | 36.08 | 4.95 | 7.8 | 1.04 |
| 13 | stress A | Stress | 13.57 | 1.08 | 58.83 | 3.81 | 7.5 | 0.90 |
| 76 | stress C | Stress | 14.40 | 2.14 | 8.32 | 3.91 | 7.8 | 0.82 |
| 65 | stress C | Stress | 15.30 | 3.52 | 15.31 | 4.89 | 8.1 | 0.92 |
| 40 | stress B | Stress | 17.21 | 7.9 | 51.67 | 3.39 | 7.2 | 0.91 |
| 46 | stress B | Stress | 17.62 | 1.95 | 17.71 | 4.14 | 7.5 | 0.98 |
| 72 | stress C | Stress | 17.67 | 4.22 | 12.18 | 3.96 | 7.5 | 0.94 |
| 75 | stress C | Stress | 19.75 | 4.75 | 16.39 | 5.65 | 8.5 | 0.92 |
| 50 | stress B | Stress | 21.26 | 9.45 | 46.36 | 3.38 | 7.3 | 0.87 |
| 41 | stress B | Stress | 22.15 | 6.29 | 63.40 | 6.23 | 8.4 | 1.05 |
| 77 | stress C | Stress | 25.39 | 1.31 | 12.16 | 3.51 | 7.4 | 0.87 |
| 47 | stress B | Stress | 28.51 | 2.6 | 39.36 | 3.83 | 7.4 | 0.94 |
| 35 | stress B | Stress | 28.70 | 1.01 | 65.81 | 4.30 | 8.0 | 0.84 |
| 17 | stress A | Stress | 29.10 | 4.87 | 33.42 | 5.23 | 8.4 | 0.88 |
